# A Systems Immunology Approach Identifies Cytokine-Induced STAT Signaling Pathways Critical to Rheumatoid Arthritis Disease Activity and Treatment Response

**DOI:** 10.1101/691865

**Authors:** Jason Ptacek, Rachael E. Hawtin, Dongmei Sun, Brent Louie, Erik Evensen, Barbara Mittleman, Alessandra Cesano, Guy Cavet, Clifton O. Bingham, Stacey S. Cofield, Jeffrey R. Curtis, Maria I. Danila, Chander Raman, Richard Furie, Mark C. Genovese, William H. Robinson, Marc C. Levesque, Larry W. Moreland, Peter A. Nigrovic, Nancy A. Shadick, James R. O’Dell, Geoffrey M. Thiele, E. William St Clair, Christopher C. Striebich, Matthew B. Hale, Houman Khalili, Franak Batliwalla, Cynthia Aranow, Meggan Mackay, Betty Diamond, Garry P. Nolan, Peter K. Gregersen, S. Louis Bridges

## Abstract

Rheumatoid arthritis (RA), a chronic autoimmune disease characterized by circulating autoantibodies, involves many cytokine-mediated signaling pathways in multiple immune cell subsets. Most studies of immune cells in RA have limitations, such as analysis of a small number of cell subsets or pathways, and limited longitudinal data on patient phenotypes. In this study, we used an innovative systems immunology approach to simultaneously quantify up to 882 signaling nodes (Jak/STAT signaling readouts modulated by cytokines and other stimuli) in 21 immune cell subsets. We studied 194 RA patients and 41 controls, including 146 well-characterized RA patients prior to, and 6 months after, initiation of methotrexate or biologic agents from the Treatment Efficacy and Toxicity in RA Database and Repository (TETRAD). There was strikingly attenuated signaling capacity in RA patients in IFNα stimulation followed by measurement of phosphorylated STAT1 [IFNα→p-STAT1] in six immune cell subsets. Multiple nodes showed negative association with disease activity, including IFNα→STAT5 signaling in naive and memory B cells. In contrast, IL-6-induced STAT1 and STAT3 activation in central memory CD4-negative T cells showed a positive association with disease activity. Multiple nodes were associated with treatment response, including IFNα→STAT1 in monocytes and IL-6→STAT3 in CD4+ naive T cells. Decision tree analysis identified a model combining these two nodes as a high-performing classifier of treatment response to TNF inhibitors. Our study provides novel information on RA disease mechanisms and serves as a framework for the discovery and validation of biomarkers of treatment response in RA.

## Introduction

Rheumatoid arthritis (RA) is a heterogeneous, chronic systemic disease characterized by synovial inflammation, cartilage destruction, and progressive bony erosions, often leading to joint damage and disability (*1*). Although its cause is unknown, RA is characterized by dysregulation of immune responses through production of autoantibodies and cytokines. T lymphocytes play a key role in RA, as supported by the strong HLA associations (*2-4*) as well as more recent identified genetic associations that indicate the involvement of T cell pathways (*5, 6*). However, given the cellular diversity within the inflamed joints, including dendritic cells, monocytes, macrophages, mast cells and B cells, many cell types (and the interactions between them) are now known to play essential roles in RA pathogenesis (*7*). Dysregulation of cytokines such as TNF and interferons, and intracellular signaling pathways in different cell subsets that stimulate Jak/STAT pathways among others, are thought to mediate the chronic inflammatory response in RA (*8-11*).

Investigations into the pathologic roles of peripheral blood immune cells in RA have typically focused on one or a few cell types, as analyzing multiple subsets of cells from relatively small volumes of blood is challenging. Single-cell network profiling (SCNP) is a systems-level, multi-parametric flow cytometry technique that simultaneously interrogates intracellular signaling pathways in multiple immune cell subsets within heterogeneous populations such as peripheral blood mononuclear cells (PBMCs) (*12, 13*). This approach allows measurement of both basal (unstimulated) intracellular cell signaling and changes induced by stimulation of cells with specific phenotypes of interest. The combination of a stimulus (e.g. IFNα) and the associated intracellular readout [such as phosphorylation of STAT1 (p-STAT1)] can be assessed simultaneously in multiple cell types in a single well. This technique has been applied to characterize immune responses in healthy subjects (*14*), to identify prognostic biomarkers in chronic lymphocytic leukemia (*15*) and to validate predictors of response to induction therapy in acute myeloid leukemia (*16, 17*).

The treatment of RA could be significantly improved by the development of reliable prognostic and predictive biomarkers that can be applied at the level of the individual. This would include biomarkers of disease activity in particular pathways as a given time, as well as biomarkers of response to treatment by measuring changes in responders vs non-responders to a particular drug. This precision medicine approach is particularly important in RA as there are multiple FDA-approved drugs directed against a variety of targets, including TNF (etanercept, infliximab, adalimumab, golimumab, certolizumab), IL-1 (anakinra), IL-6 receptor (tocilizumab, sarilumab), T cells (abatacept), B cells (rituximab), and Janus kinases (tofacitinib, barictinib)(*1*). In addition, there are many investigational new drugs being developed, and biosimilar drugs are being increasingly used (*18*). Lack of appropriate biomarkers may lead to suboptimal treatment responses that may lead to structural joint damage, organ damage (e.g. lung), diminished physical and social function, and inability to participate in the workforce. Stratifying patients based on pathogenic pathways may improve the efficiency of clinical trials of investigational new drugs. Additional potential applications of biomarkers include identification of RA patients at high risk of joint damage, patients at risk of developing organ system involvement such as interstitial lung disease and monitoring for impending loss of therapeutic effectiveness.

In this study, we address the important issues of identifying cell signaling patterns in cell subsets that may be associated with disease activity in individual patients, and how these patterns change before and after treatment with methotrexate (MTX) or biologic agents. These data will contribute to better targeted therapy in RA. For example, subjects who have active disease characterized by upregulated STAT signaling in response to IL-6 may respond better to an IL-6 inhibitor than a drug directed at another target. Similarly, examining pre- and post-treatment profiles may lead to additional insights that will improve clinicians’ choice of therapeutic agents. After validating the reproducibility of these assays over time in healthy controls (with longitudinal samples), we quantified signaling in PBMCs from well characterized RA patients and healthy controls to gain insight into the pathogenesis of RA and its disease activity. In addition, we gleaned insights into changes induced by treatment in RA using prospectively collected clinical data of biological samples collected before and 6 months after starting treatment with methotrexate or biologic agents from the NIH-funded Treatment Efficacy and Toxicity in RA Database (TETRAD) study.

We demonstrate several important findings. Immune cells from RA patients generally showed reduced response to stimuli. More specifically, IFNα induced STAT signaling, and other specific signaling nodes, are significantly lower in immune cells from RA patients compared to controls. Monocytes from RA patients show attenuation of signaling in 7 nodes, particularly the GM-CSF+IL-2→p-STAT5 node, and some nodes show bimodal responses. RA disease activity was associated with IFNα→p-STAT5, IL-10→p-STAT1 signaling and three TCR signaling nodes. After treatment, some signaling nodes (e.g. IFNα→p-STAT5, IL-10→p-STAT1 and GM-CSF + IL-2→p-STAT5) increased in monocytes, B cells and some of the cytotoxic T cell subsets. IL-6/IL-10→p-STAT3 signaling decreased in helper T cell subsets following initiation of RA therapy. In particular, patients with higher IL-6 signaling in naive CD4+ T cells and lower IFNα signaling in monocytes showed greater therapeutic response to TNFi over time. Our study provides substantial insights into the heterogeneity of the cellular and molecular pathogenesis of RA. This approach has great potential to stratify patients according to likelihood of response to particular therapies, thus adding greatly to precision medicine in RA.

## Results

### Characterization of variability and longitudinal stability of findings in healthy controls

To determine the stability of intracellular signaling across multiple immune cell subsets, we analyzed serial samples of PBMC from healthy controls (n=11) at two time points at least one month apart. A total 27 nodes in 18 cell subsets were compared between basal and stimuli state (Supplemental Figure 1). Linear regression analysis of signaling (log2Fold) measured in longitudinal samples showed highly consistent responses over time, with correlation r^*2*^ values approaching 1 (Supplemental Figure 1A, B, and C).

Signaling stability was evaluated through calculating the ratio of the signaling variability between time points to the mean signaling level. Of 258 node:subset combinations (Supplemental Figure 1E),160 (62%) had a variability ratio <0.1 (i.e. the variability between time points was <10% of the average signaling); 227 (88%) had a ratio <0.2. Naive CD4-negative (effector) T cells had the least variability in signaling between the two time points (median ratio 0.066, range 0.03 – 0.13). Memory B cells had the most variability, although this was still modest (median ratio 0.15, range 0.04 – 0.43). Taken together, signaling capacity, as measured by SCNP, is a stable, robust phenotype over time in healthy individuals, and thus can be used to characterize the pathogenesis and treatment response in patients with immune-medicated diseases such as RA.

### Characterization of RA-associated immune cell signaling

We studied RA patients enrolled by North Shore Long Island Jewish Health System (Cohort 1), RA patients from the TETRAD Study prior to initiation of methotrexate or biologic agents (TT0) and after 6 months of treatment (T6M), and healthy controls demographically matched to each RA cohort (Table 1) (*19, 20*). A set of nodes known to be relevant to the biology of RA was analyzed in each group (Figure 1).

**Table 1.** Demographic and clinical characteristics of patients of RA and healthy controls. N/A for Race/Ethnicity means not available; otherwise, not applicable.

|  |  | Cohort 1 |  |  |  | TT0 |  |  |  | T6M |  |  |  |
| --- | --- | --- | --- | --- | --- | --- | --- | --- | --- | --- | --- | --- | --- |
|  |  | Controls<br>(n=20) |  | RA<br>(n=48) |  | Controls<br>(n=10) |  | RA<br>(n=146) |  | Controls<br>(n=11) |  | RA<br>(n=98) |  |
|  |  | n | % | n | % | n | % | n | % | n | % | n | % |
| Age at Baseline |  | Median: 55.0<br>Range 42 - 80 |  | Median: 61.5<br>Range 37 - 87 |  | Median: 49.0<br>Range 40 - 60 |  | Median: 56.0<br>Range 24 - 82 |  | Median: 45.0<br>Range 29 - 60 |  | Median: 56.0<br>Range 25 - 80 |  |
| Sex | M | 6 | 30.0 | 8 | 16.7 | 4 | 40.0 | 20 | 13.7 | 4 | 36.4 | 15 | 15.3 |
|  | F | 14 | 70.0 | 40 | 83.3 | 6 | 60.0 | 126 | 86.3 | 7 | 63.6 | 83 | 84.7 |
| Race/Ethnicity | African-American | 0 | 0 | N/A | N/A | 0 | 0.0 | 16 | 10.9 | 0 | 0.0 | 8 | 8.2 |
|  | Asian | 3 | 15 | N/A | N/A | 2 | 20.0 | 8 | 5.5 | 2 | 20.0 | 6 | 6.1 |
|  | Caucasian | 10 | 50 | N/A | N/A | 5 | 50.0 | 113 | 77.4 | 5 | 50.0 | 79 | 80.6 |
|  | Other/Unknown | 7 | 35 | 48 | 100 | 3 | 30.0 | 9 | 6.2 | 3 | 30.0 | 5 | 5.1 |
| Anti-CCP | Positive |  |  | 31 | 64.6 |  |  | 91 | 62.4 |  |  | 62 | 63.3 |
|  | Negative | N/A | N/A | 17 | 35.4 | N/A | N/A | 34 | 23.2 | N/A | N/A | 22 | 22.4 |
|  | Unknown |  |  | 0 | 0.0 |  |  | 21 | 14.4 |  |  | 14 | 14.3 |
| RF | Positive |  |  | 41 | 85.4 |  |  | 103 | 70.5 |  |  | 66 | 67.4 |
|  | Negative | N/A | N/A | 7 | 14.6 | N/A | N/A | 42 | 28.8 | N/A | N/A | 31 | 31.6 |
|  | Unknown |  |  | 0 | 0.0 |  |  | 1 | 0.7 |  |  | 1 | 1.0 |
| DAS28 |  | N/A |  | Median: 3.36<br>Range 1.23 - 6.93 |  | N/A |  | Median: 4.96<br>Range 1.25 - 8.22 |  | N/A |  | Median: 3.50<br>Range 0.16 - 7.52 |  |
| Concomitant Medications | GC |  |  | 7 | 14.6 |  |  | 29 | 19.8 |  |  | 21 | 21.4 |
|  | MTX |  |  | 21 | 43.8 |  |  | 32 | 22.0 |  |  | 35 | 35.7 |
|  | GC+ MTX | N/A | N/A | 12 | 25.0 | N/A | N/A | 59 | 40.4 | N/A | N/A | 18 | 18.4 |
|  | Unknown/None |  |  | 8 | 16.7 |  |  | 26 | 17.8 |  |  | 24 | 24.5 |
| Recent Previous Biologic at Baseline | Yes |  |  |  |  |  |  | 38 | 26.0 |  |  |  |  |
|  | No | N/A | N/A | N/A | N/A | N/A | N/A | 108 | 74.0 | N/A | N/A | N/A | N/A |
| Current Biologic | Abatacept | N/A | N/A | 23 | 47.9 | N/A | N/A | N/A | N/A | N/A | N/A | 13 | 13.3 |
|  | Anti-TNF | N/A | N/A | 2 | 4.2 | N/A | N/A | N/A | N/A | N/A | N/A | 40 | 40.8 |
|  | Rituximab | N/A | N/A | 2 | 4.2 | N/A | N/A | N/A | N/A | N/A | N/A | 1 | 1.0 |
|  | Tocilizumab | N/A | N/A | 0 | 0.0 | N/A | N/A | N/A | N/A | N/A | N/A | 19 | 19.4 |
|  | None | N/A | N/A | 21 | 43.7 | N/A | N/A | 98 | 100 | N/A | N/A | 25 | 25.5 |

**Figure 1.**
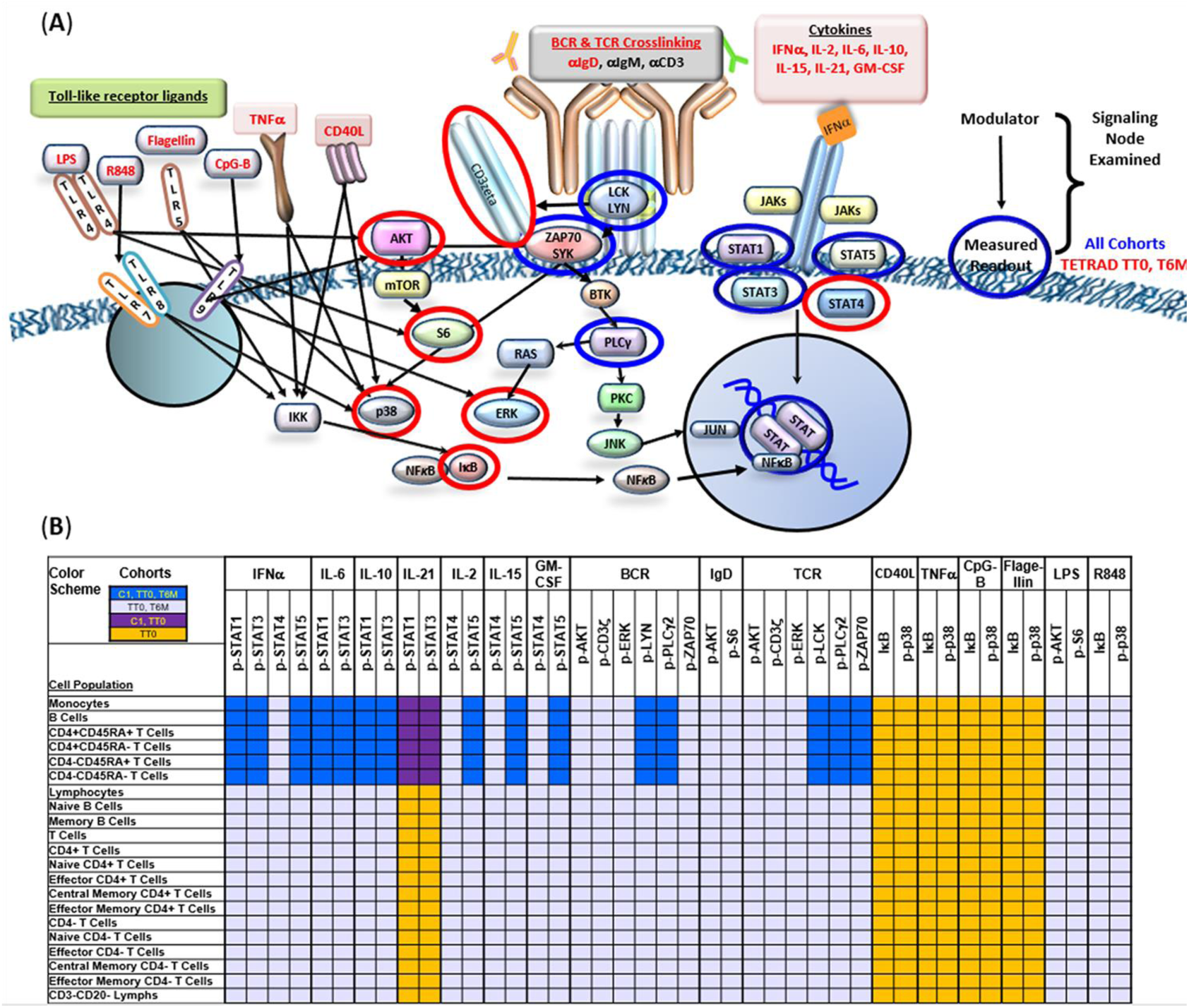
Signaling pathways and cell subsets analyzed in the study. A. In this diagram, modulators (highlighted in red) are shown outside the cell membrane and cell signaling molecules (red or blue circled) whose phosphorylation status were measured are shown inside or adjacent to the cell membrane. Signaling molecules analyzed in all cohorts are circled in blue; additional molecules analyzed in TT0 and T6M are circled in red. The term “signaling node” is used to refer to proteomic readout in the presence or absence of a specific modulator (see Methods for details). B. Summary of signaling nodes examined in cell subsets among Cohort 1, TT0, and T6M. A core set of nodes (15 in total) and cell populations (6 in total) were analyzed in all 3 sets of samples (dark blue) with additional biology analyzed in specific cohorts. In addition, due to cells availability, analyses performed in TT0 only are highlighted in yellow, analyses performed in Cohort 1 and TT0 are labeled in purple, and analyses performed in TT0 and T6M are labeled in gray. The signaling pathways of peripheral blood cells from RA patients and healthy controls were modulated using cytokines (IFNα, IL-2, IL-6, IL-10,IL-15, IL-21, GM-CSF), crosslinking antibodies to B and T cell receptors (BCR, TCR, IgD (*44*)), and TLR agonists (CD40L(*45*), TNFα, Resiquimod R848), pathogen-associated molecules (CpG-B, Flagellin and LPS) as shown on the top row. The resulting pathway activations measured are shown on the second row, and cell subsets analyzed are shown in the left column.

We initially compared modulated signaling in RA datasets to signaling in healthy controls. RA patients in Cohort 1 and TETRAD patients after treatment (T6M) all had established disease, and similarities in distribution of sex, RA disease activity (median DAS28 3.36 vs 3.50) and, importantly, treatment status (Table 1). Thus, these two RA groups were compared to controls across multiple nodes. The analysis showed a number of significant cell signaling differences between RA and controls (Figures 1, 2 and Supplemental Figure 2).

**Figure 2.**
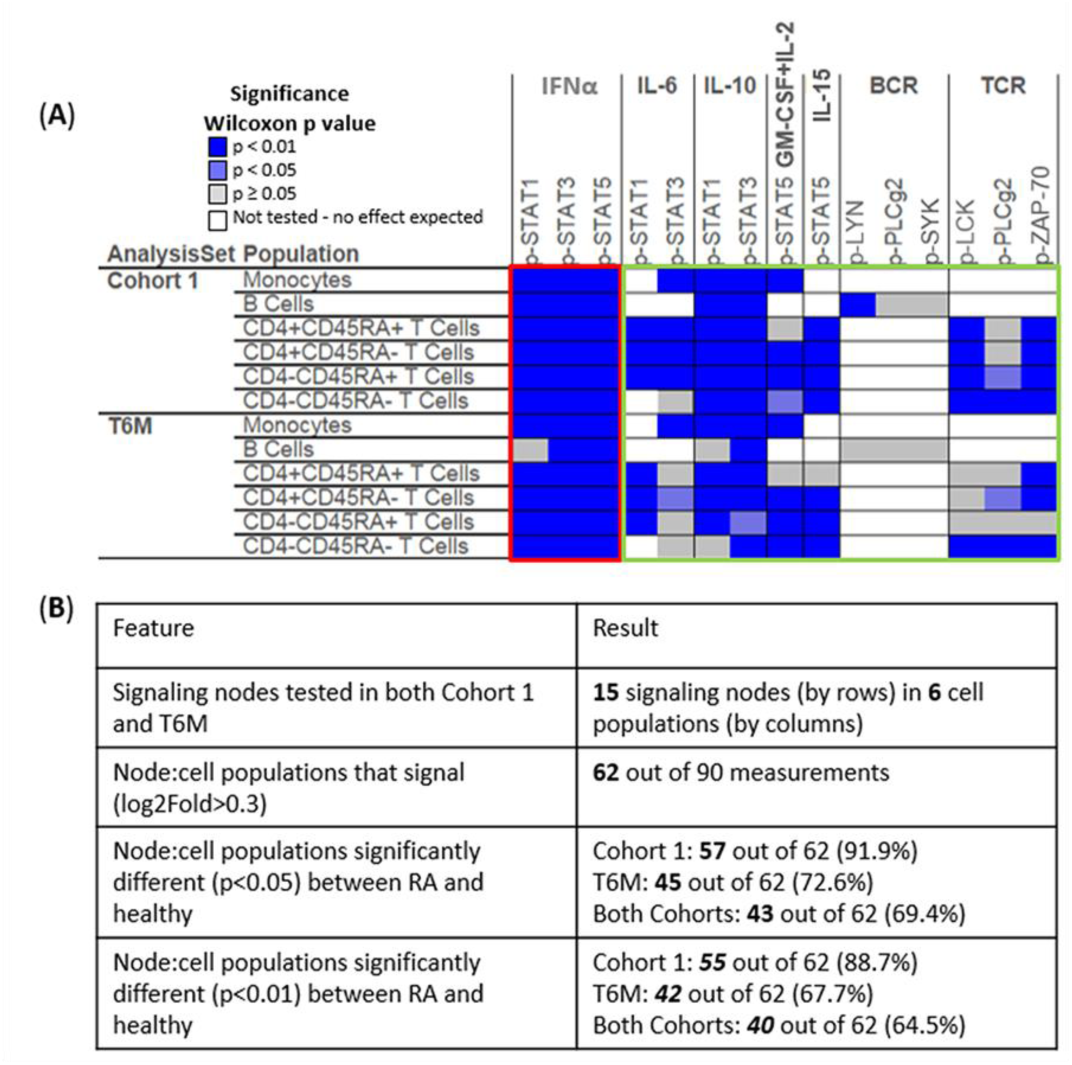
Induced signaling is significantly altered for specific nodes within immune cell subsets from patients with RA compared to healthy controls. A. Heatmap of p values of the Wilcoxon signed-rank test comparing signaling in RA compared to controls in Cohort 1 and T6M within immune cell subsets, including total monocytes, total B cells, naïve Th cells (CD4+/CD45RA+), memory/effector Th cells (CD4+/CD45RA-), naïve Tc cells (CD4-/CD45RA+), and memory/effector Tc cells (CD4-/CD45RA-). Levels of significance are shown as dark blue (p < 0.01); light blue (p < 0.05); gray (p≥ 0.05, not significant). White is used for cell populations that do not respond to the stimulus tested (for example B cell responses to TCR cross-linking). The rows of the heatmap show six populations of immune cells analyzed in both cohorts and the columns show the signaling nodes assayed. B. A total of 90 assessments of induced signaling (15 nodes in 6 cell populations) were made for both Cohort 1 and T6M.

Significant differences in induced signaling between RA immune cell subsets and controls are represented as a heatmap of Wilcoxon p values (Figure 2A), and summarized in a table (Figure 2B). There were consistent differences in IFNα induced p-STAT1, p-STAT3, and p-STAT5 signaling in monocytes, B cells, and T cell subsets in RA compared to controls, and many differences of signaling in response to IL-6, IL-10, GM-CSF + IL-2, IL-15, and TCR (Figure 2A). In T6M group, 45 out of 62 signaling nodes showed significant differences in the level of signaling compared to controls. Fifty-seven of these 62 (91.9%) also showed similar signaling differences between Cohort 1 RA patients and healthy controls. The majority of these associations also met a more stringent threshold of p value < 0.01 (Figure 2B). All these differences represented reduced signaling capacity in RA in response to the stimuli tested; none represented increased signaling capacity in RA.

A subset of the most striking and interesting findings are shown in Figure 3. In RA Cohort 1, IFNα activation of p-STAT1 was significantly lower than in controls (p < 0.01) in 6 cell subsets: total monocytes, total B cells, naïve helper Th cells (CD4+/CD45RA+), memory/effector Th cells (CD4+/CD45RA-), naïve Tc cells (CD4-/CD45RA+), and memory/effector cytotoxic Tc cells (CD4-/CD45RA-). These same findings were noted in 5 of the 6 cell subsets T6M samples compared to controls (the exception being total B cells). Of these six cell subsets, monocytes showed the most marked decreases in modulated IFNα→p-STAT1 signaling in RA (red boxes in Figure 3A, p < 0.001). As shown in Figure 3B, more detailed analysis of RA monocytes showed significantly attenuated signaling in all six nodes in both RA Cohort 1 and RA T6M. GM-CSF→p-STAT5 signaling (co-stimulated with IL-2 for T cell modulation) was the most reduced node in RA (RA Cohort 1 vs controls, p = 2.27 × 10^-21^ and RA T6M vs controls, p = 2.14 ×10^-7^) (red boxes in Figure 3B).

**Figure 3.**
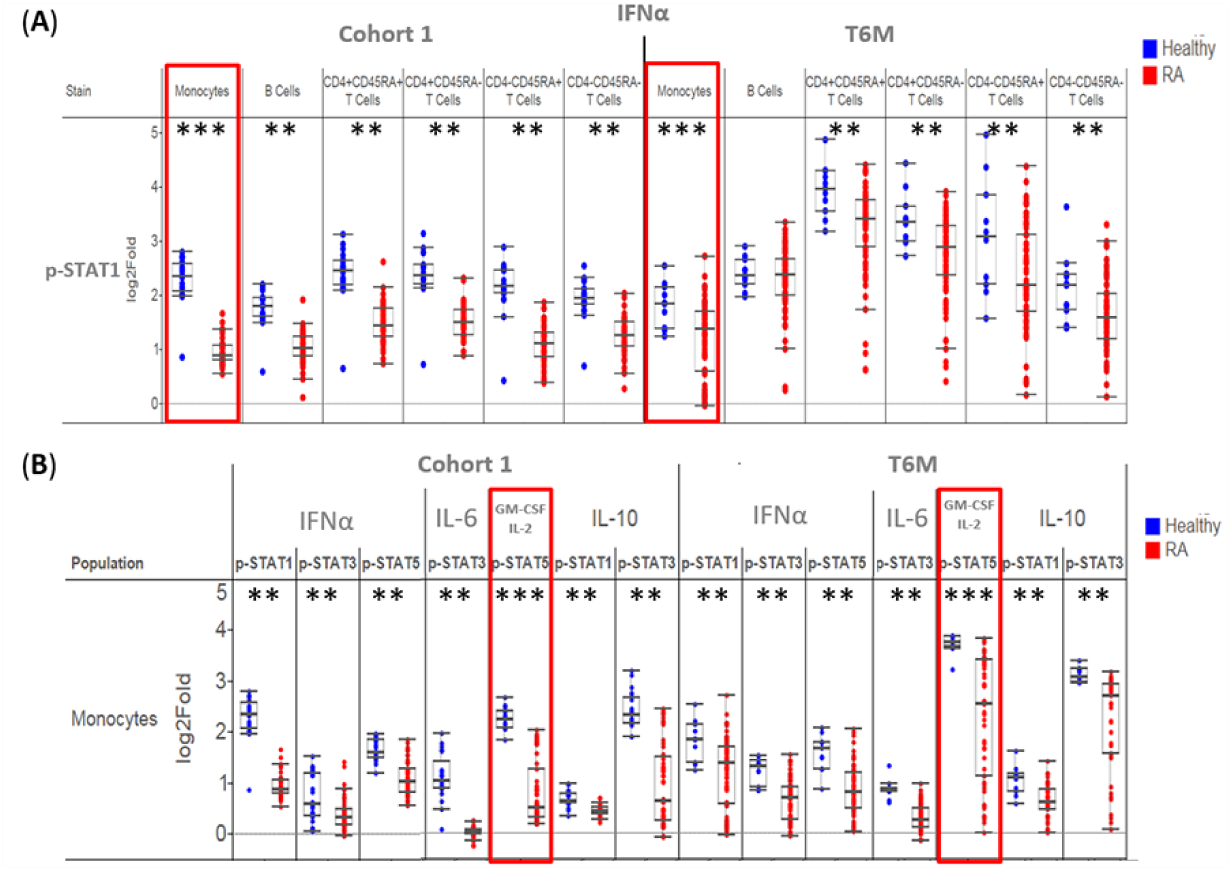
Patients with RA show consistent widespread attenuated signaling within 6 subsets of peripheral blood cells. (For the results of the full panel analyzed in these 6 cell subsets, see Supplemental Figure 2). Each dot represents one subject at one time point, with RA patients shown in red and healthy controls shown in blue. Boxes indicate the first, second (median), and third quartiles with the whiskers extending to 1.5 interquartile range (IQR). ** Differences between RA and controls were statistically significant (Wilcoxon signed-rank test) at p<0.01. *** Differences between RA and controls were statistically significant at p<0.001. A. IFNα activation of STAT1 (log2Fold) was significantly lower in RA patients from Cohort 1 (left) and T6M (right) compared to controls. B. Of the six immune populations, the signaling capacity varied the greatest in RA monocytes compared to control monocytes in both cohorts. Monocyte signaling in response to IFNα, IL-6, GM-CSF + IL-2, and IL-10 are shown with readouts of p-STAT1, p-STAT3, and p-STAT5.

In addition to characterizing changes in the overall magnitude of signaling within a defined cell population, we were able to quantify the proportion of cells within a cell subset that are responding to modulation. Figure 4A shows an example from one healthy control and one representative RA patient in which we examined basal cell signaling in monocytes (left panels), and modulated cell signaling after stimulation with IFNα (center panels) and GM-CSF (right panels), using phosphorylated STAT5 as a readout. Compared to the control, monocytes from this RA patient showed a bimodal signaling response, with some cells responding and others not, in the GM-CSF→STAT5 node. Figure 4B shows the striking differences between RA patients and healthy controls. Among RA patients in Cohort 1, the median proportion of monocytes responding to GM-CSF in the STAT5 pathway was 36% with a high degree of variability (range 5-83%) compared to 90% in controls (range 79-96%). Similar findings were seen in the T6M dataset (median RA response 57%, range 2-95%). A similar diminished and bimodal response was seen in the IL-10→STAT3 node (data not shown); no other nodes in monocytes with reduced signaling in RA showed this bimodal response.

**Figure 4.**
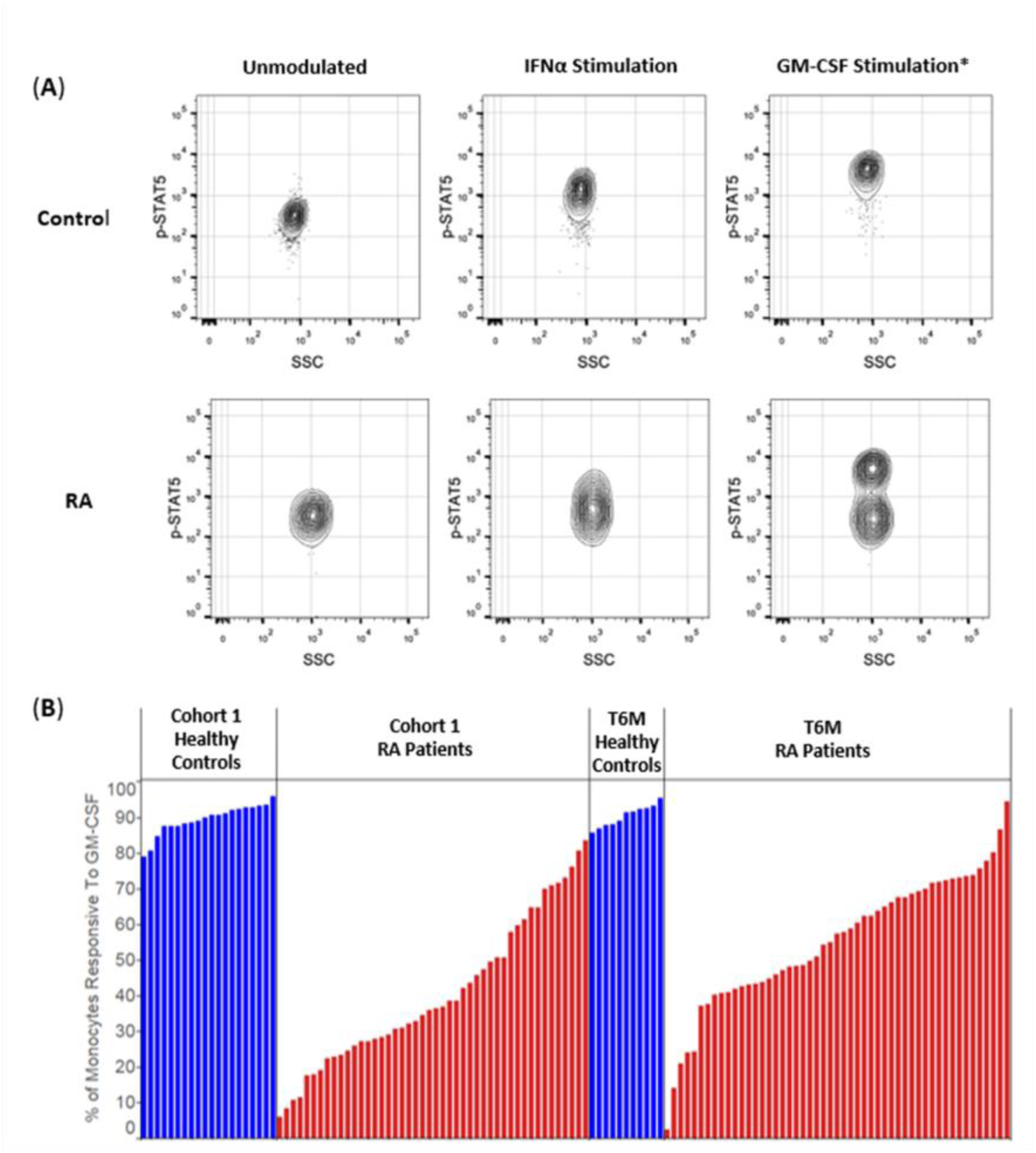
Monocyte Jak/STAT signaling showed bimodal response to GM-CSF in RA compared to healthy controls. A. Representative contour plots show p-STAT5 in monocytes from one healthy control and from one RA patient under three different conditions: basal (unmodulated); IFNα stimulation, and GM-CSF + IL-2 stimulation. Monocytes from the RA patients showed a bimodal GM-CSF→p-STAT5 response whereas IFNα→p-STAT5 was unimodal (*Note: IL-2 was included to induce STAT5 signaling in T cells; STAT5 signaling is induced in monocytes from GM-CSF alone. To assess more nodes with limited patient material, cytokines with non-overlapping cell specificity were profiled together. In this case IL-2 with T cell activating characteristics and GM-CSF with specificity to monocytes were combined.) B. Histograms show percentages of monocytes that respond to GM-CSF from RA patients and controls. Monocytes from RA patients of Cohort 1 and T6M, but not from controls, showed a bimodal GM-CSF→p-STAT5 response.

### Identification of signaling patterns associated with RA disease activity

To evaluate the association of signaling characteristics with RA disease activity (as assessed by DAS28 scores), we applied linear regression analysis (including age as a covariate) to the TT0 dataset. The TT0 dataset comprised 146 patients about to begin therapy with MTX or biologic agents for clinical indications. These patients had higher disease activity (median DAS28 4.92) than Cohort 1 (median DAS28 3.36) and as expected, had lower disease activity after treatment for 6 months (T6M) (median DAS28 3.50) (Table 1). Multiple nodes showed evidence of association with disease activity (p<0.05, adjusting for age), being either inversely correlated (Figure 5A, blue heatmap) or, less frequently, positively correlated (Figure 5A, red heatmap). Representative examples are shown in Figures 5B-C. The IFNα→p-STAT5 node was associated with disease activity (p<0.05) in 12 different PBMC subsets (red box, Figure 5A). Similarly, IL-10→p-STAT1 was associated with disease activity in 8 cell subsets (black box, Figure 5A). Signaling through p-Lck, p-CD3-ζ and other readouts was associated with disease activity in multiple T cell subsets (purple box, Figure 5A). Higher disease activity was associated with decreased modulated signaling in some nodes including IFNα→p-STAT5 in CD4-/CD45RA+ T cells (Figure 5B) and with increased modulated signaling in other nodes, for example IL-6→p-STAT3 in central memory CD4-T cells (Figure 5C).

**Figure 5.**
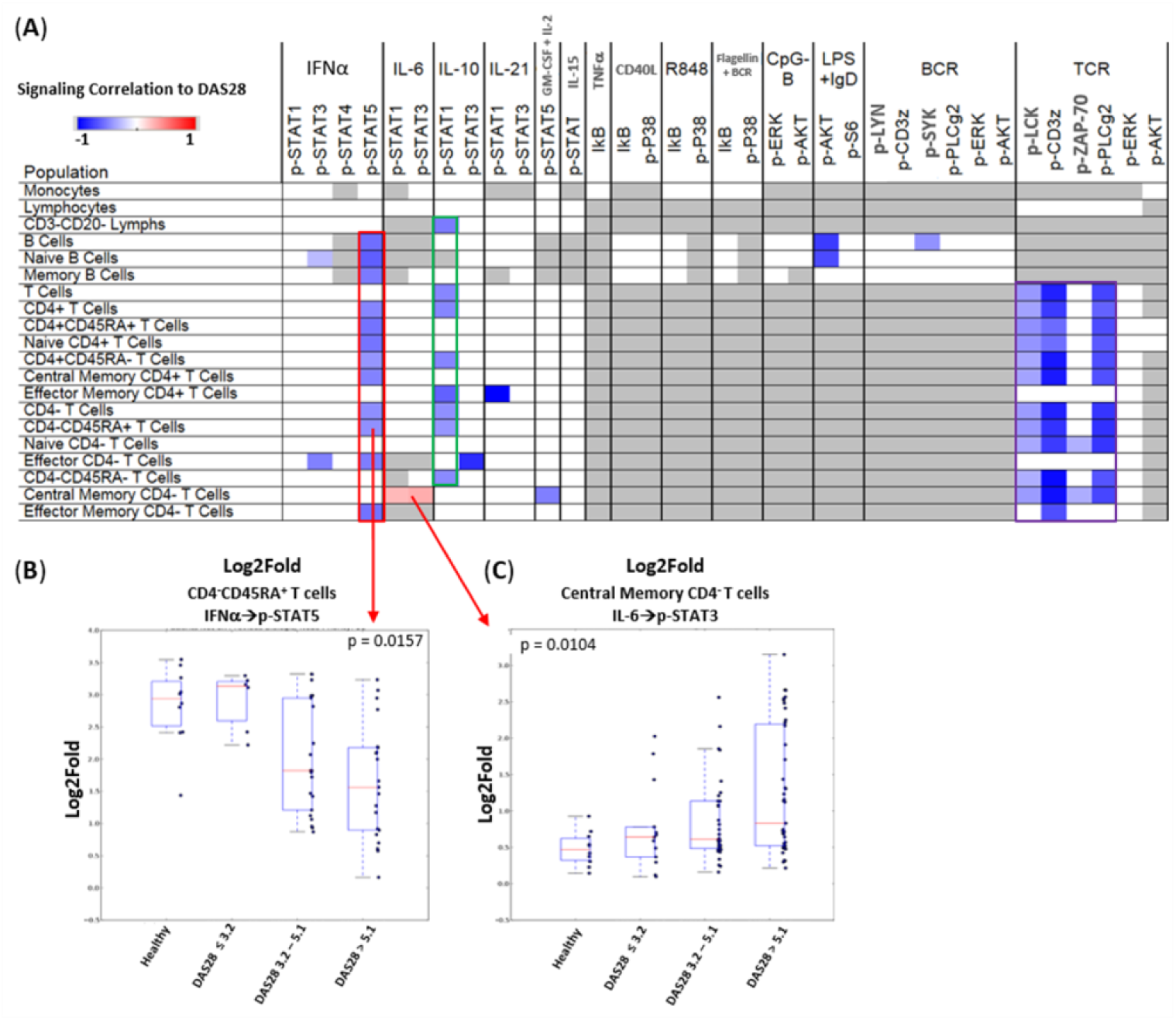
Analysis of signaling associated with RA disease activity (DAS28). A. The heatmap shows the Spearman’s correlation (r) between signaling and DAS28 among TT0 patients that were not on a biologic agent, organized by row with the 20 immune cell populations and by column with the signaling nodes assayed. Signaling nodes in specific cell populations that were significantly (p < 0.05, adjusting for age) and positively associated with disease activity are colored in red. Nodes that inversely correlated with disease activity are colored in blue, and nodes that were not significantly correlated with disease activity are colored in white. Nodes that do not signal in specific cell population are colored in gray. (B – C). Representative signaling nodes associated with disease activity in TT0. Analyses were performed by logistic regression, adjusting for patient age. Data are represented as Log2Fold metric stratified by disease activity categories. Boxes indicate the first, second (median), and third quantiles with the whiskers extending to 1.5 interquartile range (IQR).

### Analyses pre- and post-treatment reveals both stable and dynamic signaling nodes and facilitates development of classifiers of treatment response

Paired analyses were performed in T6M (84 patients) and TT0 for 32 nodes in 21 immune cell populations (Figure 1) before and after treatment. Figure 6 shows nodes and cell populations in which signaling significantly changed or remained stable from baseline to 6 months following initiation of treatment. Nodes that demonstrated increased responsiveness at T6M include IFNα→p-STAT5 in monocytes, B and cytotoxic T cell subsets and GM-CSF + IL-2→p-STAT5 in cytotoxic T cell subsets (black boxes). Lymphocytes, which include T, B, and NK cell subsets, became less responsive in the IL-6→p-STAT1 node. Interestingly, whereas IL-6→p-STAT3 and IL-10→p-STAT3 responses were diminished in helper T cell subsets (red boxes), IL-10 induced statistically significant greater p-STAT1 signaling in monocytes, B and T cell subsets (purple box). IFNα→p-STAT1 and IL-6→p-STAT1 were relatively stable before and after treatment. BCR signaling was higher in naïve B cells and lower in memory B cells after treatment whereas TCR signaling was generally weak.

**Figure 6.**
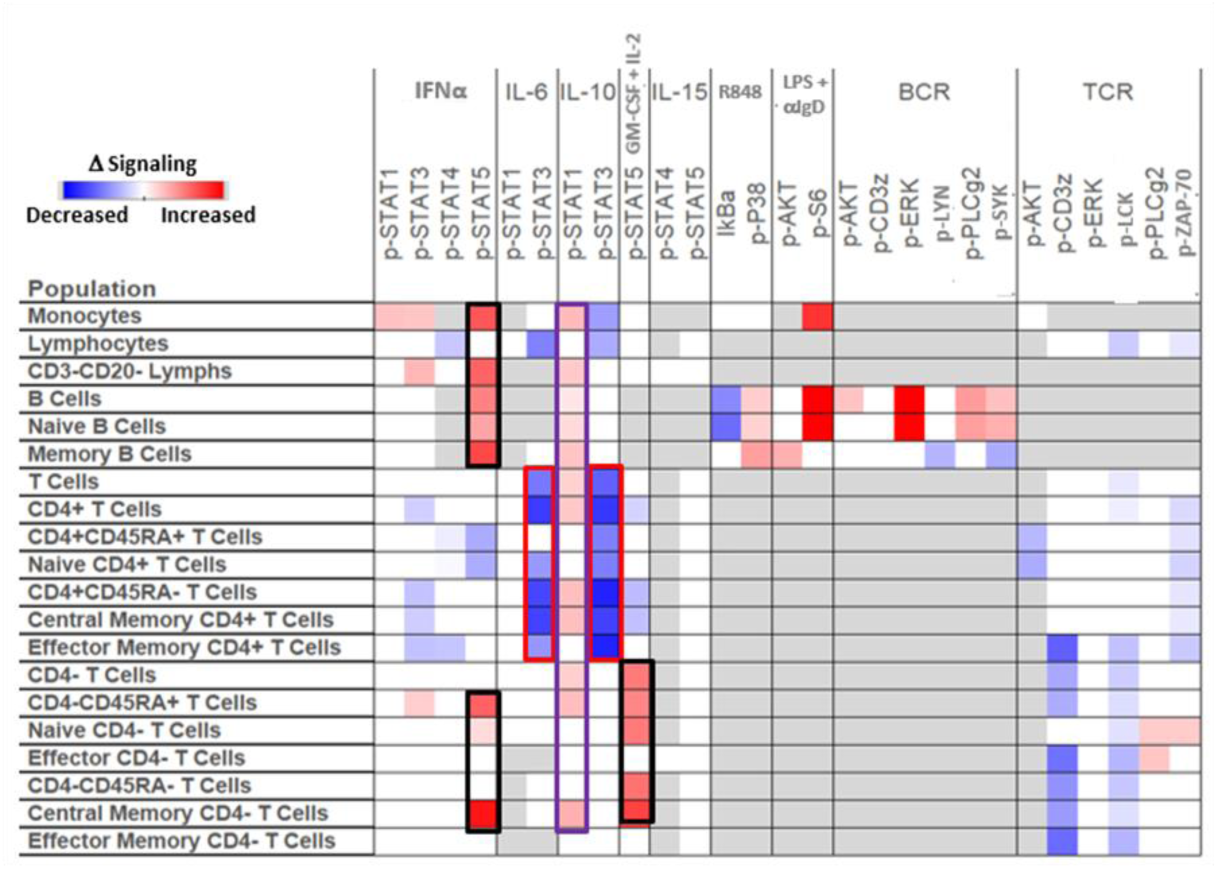
Longitudinal analysis of signaling from TETRAD RA patients at TT0 and T6M. The heatmap shows by row the 20 immune cell subsets and by column the signaling nodes assayed. Signaling nodes in specific cell populations that significantly (Wilcoxon signed-rank test) increased in their modulated response at 6 months compared to baseline are colored in red, nodes with significantly reduced signaling are colored in blue, and nodes that show similar levels of signaling between time points are colored in white. Nodes that do not signal in the specific cell population are colored in gray.

We performed analyses to identify patients with good vs poor response to TNFi based on the association between signaling and treatment response. Decision tree analysis and Least Absolute Shrinkage and Selection Operator (LASSO) regression analysis were employed for these hypothesis-generating analyses. Pre-treatment (TT0) and post-treatment (T6M) SCNP data from 46 RA patients not previously on a biologic agent and initiating TNFi therapy (23 etanercept, 17 adalimumab, 4 infliximab, 2 golimumab) were analyzed (see Supplemental Table 1 for characteristics of these patients). Of these 46 subjects, 9 (20%) met criteria for good EULAR response at 3 months, 20 (43%) had moderate response, and 17 (37%) had none. Decision tree analysis identified models that each combined two nodes to classify response/nonresponse to TNFi (Figure 7A). Among our top-performing classifiers was a model combining IFNα→p-STAT1 signaling in monocytes with IL-6→p-STAT3 in naive CD4+ T cells, which showed that responders tended to cluster together, exhibiting higher IL-6 signaling. Non-responders showed greater signaling heterogeneity with some having high IFNα signaling or low IL-6 signaling, suggesting that there may be multiple signaling phenotypes associated with lack of response to TNFi treatment.

**Figure 7.**
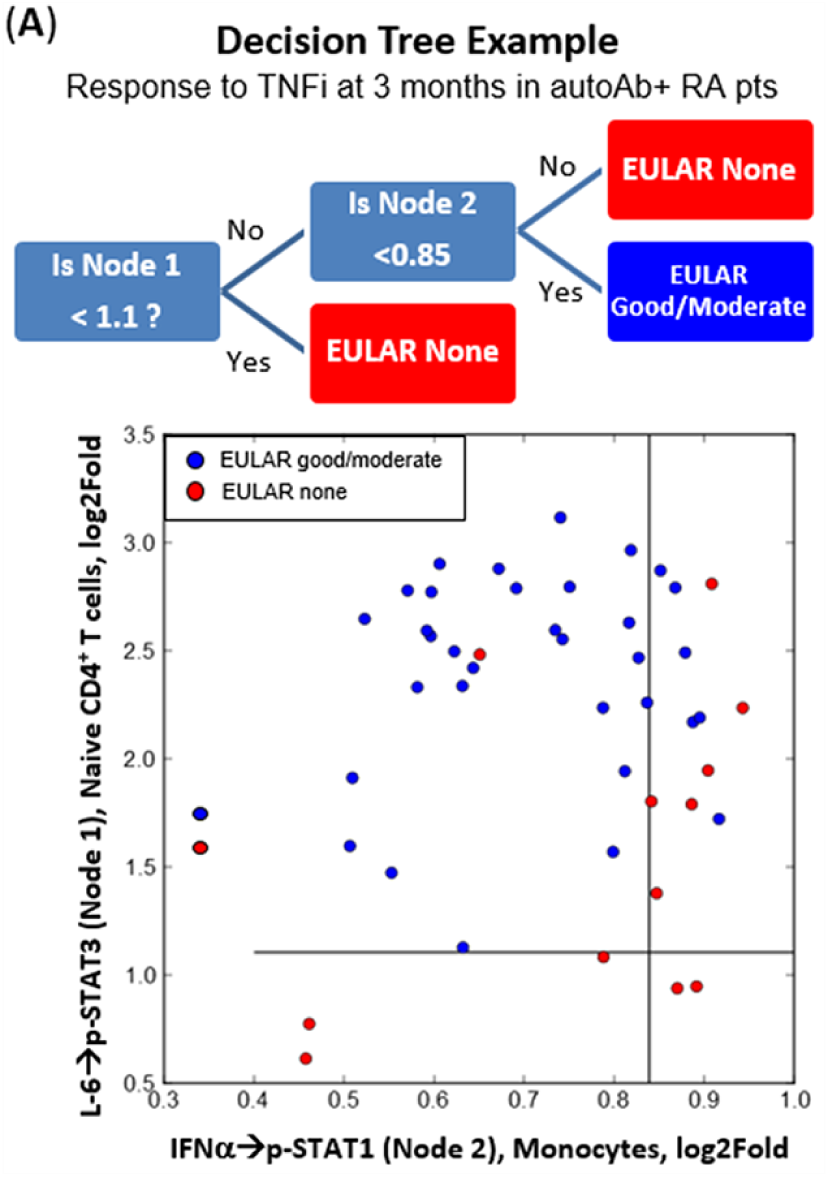
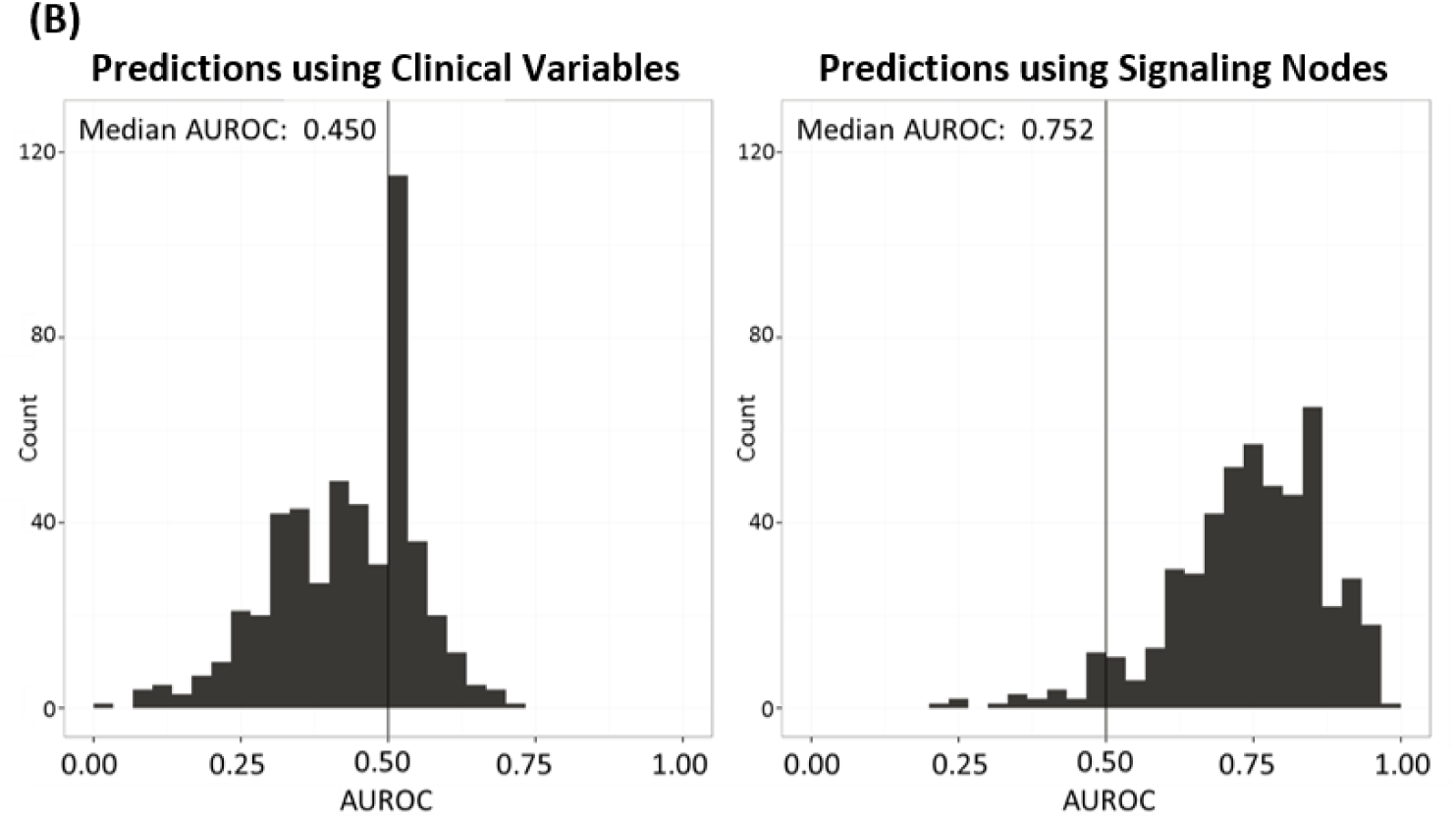
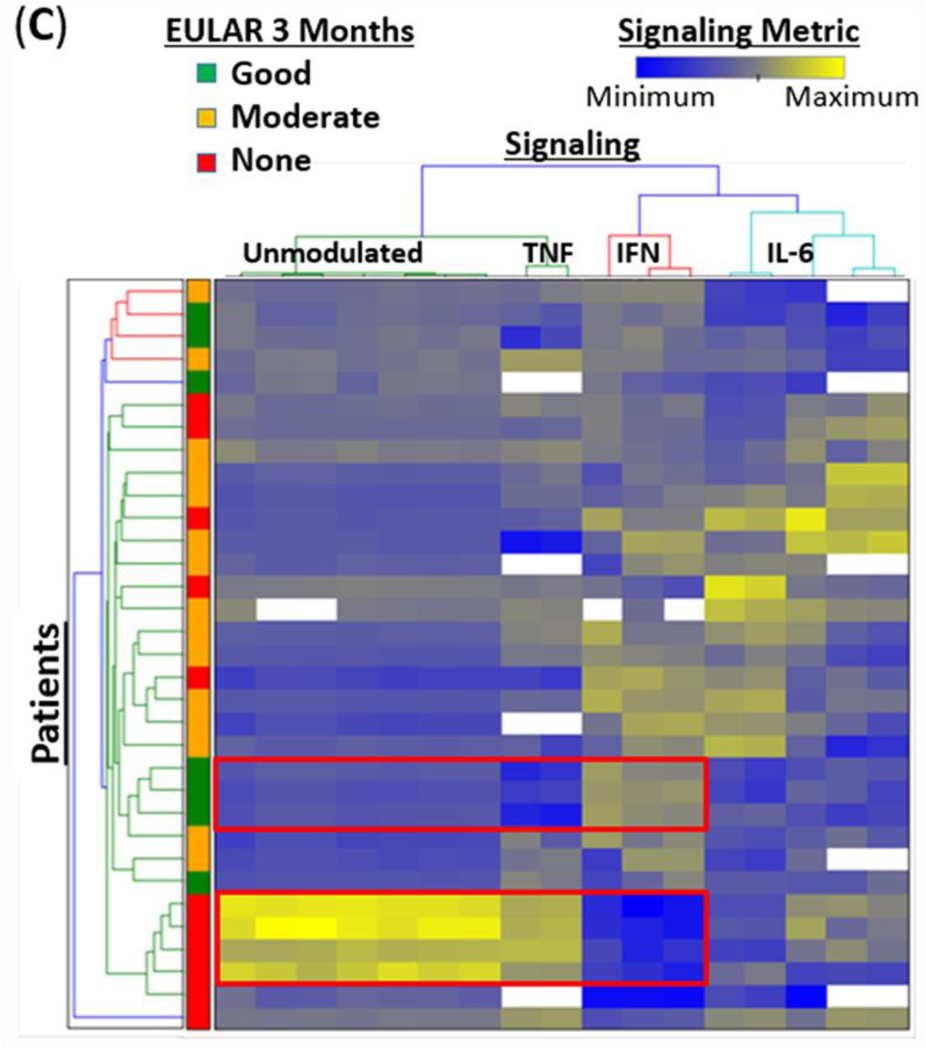
SCNP reveals functional differences between EULAR response categories in 46 patients beginning TNF inhibitors (TNFi) and can be used to develop hypothesis generating classifiers of treatment response. Of these 46 subjects, 9 (20%) met criteria for good EULAR response at 3 months, 20 (43%) had moderate response, and 17 (37%) had none. A. The combined pattern of IL-6→p-STAT3 signaling in naive CD4+ T cells and IFNα→p-STAT1 in monocytes shows a trend toward separation of EULAR responders (good or moderate) from non-responders. B. The range of performance observed from over 500 bootstraps of multivariate models using either clinical variables or signaling metrics. This example shows that the AUROC is more favorable for signaling nodes (Median AUROC=0.752) than for clinical variables (Median AUROC=0.450), suggesting it may provide a better predictor of treatment response in this exploratory cohort. C. Unsupervised clustering analysis of nodes that had univariate associations to TNFi response controlling for age and baseline DAS28 in autoantibody positive RA patients (TT0). Patients are classified as having “Good”, “Moderate” or “None” based on EULAR response criteria. Signaling metrics for the unmodulated state and nodes associated with 10 broad biologic pathways are defined: 2 TNF-related, 3 IFNα-related, and 5 IL-6-related nodes, shown as maximal signaling (yellow) or minimal signaling (blue). Red boxes highlight areas exemplifying similar and opposite patterns of unmodulated, TNF and IFN stimulated signaling between none and good responders in the TNFi treated patients.

We also applied the LASSO regression method to evaluate prediction models for EULAR response using either the collection of signaling data (56 nodes across cell subsets) or the clinical data, controlling for baseline DAS28, age, autoantibody positivity, etc. We characterized the performance of these models across 500 bootstraps samples of the data. Figure 7B shows the histograms of area under the receiver operating curves (AUROCs) for models based on clinical data and models built from signaling responses. The median AUROC for clinical models was 0.450 compared to 0.752 for the signaling models, indicating that immune cell signaling may serve as a better marker of treatment response than clinical variables. Figure 7C shows results of unsupervised clustering analysis of RA patients with nodes that had univariate associations to TNFi response controlling for age and baseline DAS28. Although there is considerable variability in signaling nodes among patients in different categories of treatment response, there are some nodes that are consistent among these categories. As highlighted in the heatmap (red boxes), non-responders tended to have elevated unmodulated signaling, moderate TNF-related and minimum IFN-related signaling, while good responders tended to show the opposite patterns.

## Discussion

We performed a comprehensive analysis of basal and modulated signaling capacity in PBMC samples from RA patients and healthy controls from two independent cohorts, among RA patients with different levels of disease activity, and before and after initiation of treatment with MTX or biologic agents. Intracellular signaling is stable over time in most subsets of PBMCs in healthy individuals, but there are consistent differences between RA patients and controls. In addition, there are patterns associated with RA disease activity. Pre- and post-treatment analysis revealed both stable and dynamic signaling nodes, and changes associated with response or lack of response to treatment, especially TNFi.

Many of the nodes that differ in RA compared to controls involve IFNα as a modulator of cell signaling. IFNα has pleiotropic effects, including the promotion of proliferation and clonal expansion of T cells (*21*), increasing antibody production by B cells (*22*), and with enhancing innate immune cell function. An interferon signature, previously associated with active or severe disease in SLE (*23*), has been implicated in other autoimmune diseases such as Sjögren’s syndrome (*24, 25*). Involvement of interferons in the pathogenesis of RA and possibly in therapeutic response has been reported (*26, 27*). We have previously reported that among TETRAD subjects, increased pretreatment serum IFN-β/α ratio is strongly associated with non-response to TNFi (*28*). Our observation that IFNα signaling differs between RA and healthy controls across monocytes, B cells and T cell subsets (Figure 2 and Figure 3) adds to this body of evidence. Reduced signaling capacity in the IFNα→p-STAT5 signaling node in several cell subsets is associated with disease activity (Figure 5) and significantly increased after six months of treatment (Figure 6), suggesting that signaling capacity in this node increases across multiple immune cell subsets under therapeutic pressure. Determination of whether this change is associated with clinical benefit will require further studies.

Interestingly, IL-10→p-STAT1 signaling in B and T cells was also associated with disease activity (Figure 5). IL-10 is another key regulator of the inflammatory response and can activate STAT1 and STAT3 through activation of JAK1 and TYK2. The major role of IL-10 is to block the antigen specific T-cell cytokine response. IL-10 co-stimulation with CD3 and CD28 produce more IFNγ in peripheral blood CD4+ T cells from patients with RA than in CD4+ T cells from healthy controls (*29*). IL-10 can also inhibit TNFα production in lipopolysaccharide-stimulated macrophages (*30*). The opposing directionality (Figure 6) of IL-10→p-STAT1 (increasing after treatment with TNFi) and IL-10→p-STAT3 (decreasing after treatment with TNFi) suggest different roles of STAT1 and STAT3 in the IL-10 signaling pathway. The negative association of disease activity with increased IL-10→p-STAT1 (Figure 5) and the increased IL-10→p-STAT1 signaling observed after initiation of TNFi treatment (Figure 6) indicate that enhanced JAK1/STAT1 signaling may be beneficial in RA. Furthermore, it suggests that in addition to the direct inhibition of TNF production, TNFi may also improve RA through its effects on IL-10 signaling. Further analysis of the possible clinical benefit of increase IL-10→p-STAT1 signaling during TNFi is warranted.

Several interesting differences in monocyte signaling were identified between RA and healthy controls. The observation that there is decreased responsiveness of monocytes to stimulation with GM-CSF, and the bimodal response in this cell population (Figure 3 and 4), are of particular interest given the initial success of anti-GM-CSF mavrilimumab in clinical trials of RA (*31*). GM-CSF is an inflammatory cytokine with strong activities cross myeloid cells populations, including DCs, macrophages, and monocytes. GM-CSF deficiency results in impaired phagocytosis and GM-CSF deficient mice develop an SLE-like phenotype (*32*). In RA, it is possible that the substantial number of monocytes refractory to GM-CSF may contribute to reduced clearance of apoptotic cells, contributing to inflammation. Another possible explanation is that reduced GM-CSF signaling in RA results from chronic exposure of monocytes to GM-CSF produced to differentiate monocytes into an M1 inflammatory macrophage subset (*33*). The bimodal response identified in our analysis may reflect the diversity of cells within the pool of CD14+ monocytes, with exhausted cells comprising the non-responsive population. It is possible that inhibition of GM-CSF signaling in RA removes the chronic stimulus and facilitates a “re-setting” of the responsiveness of this population, leading to a bimodal distribution.

Combining the responses to multiple disease-relevant stimuli in multiple immune cell types allows changes, both large and small, in the overall immune signaling architecture to be assessed for their predictive clinical power. The use of multiparametric datasets (Figure 7) may thus enable the development of algorithms to predict treatment response in RA. In our analysis, a decision tree combining IL-6 and IFNα signaling responses identifies responders to TNFi treatment better than either node alone. Our data suggest that multiparametric signaling data is more strongly associated with TNFi with response than are clinical variables alone. Critical to confirmation of any such analysis will be access to adequately sized and clinically well-annotated training and validation cohorts.

Our findings provide valuable insights into the precise immunologic mechanisms driving RA disease activity in individual patients. These insights could lead to improved understanding of mechanisms of disease activity and treatment response. Such understanding would apply not only to TNFi, but inhibitors directly targeting the JAK signaling pathway (tofacitinib, baricitinib, upadacitinib, filgotinib) and also methotrexate and tocilizumab, which have recently been reported to have effects on the JAK/STAT pathway (*34, 35*).

One perceived weakness of our study is that we performed analyses on immune cells from peripheral blood rather than from synovial tissue. Many important insights have been gained by analysis of cells from RA synovial tissue (*36*), but such samples are difficult to obtain outside of research studies. Furthermore, compared to analysis of peripheral blood cells, analysis of synovial tissue is not feasible as a biomarker of disease activity or treatment response. It is also important to note that in order to determine the generalizability of these findings, additional populations (e.g. those of African or Asian ancestry; autoantibody negative; early, treatment-naïve RA, etc.) will need to be studied.

In summary, we have used a systems immunology approach to cell signaling in subsets of peripheral blood mononuclear cells in well-characterized patients with RA. The important biologic insights provided by these studies will serve as the basis for future research to better understand the pathobiology of RA, immunologic correlates of important endo-phenotypes such as autoantibody positivity (*10*), mechanisms of disease activity, and to inform future clinical trials by allowing optimization of patient selection. Furthermore, these data will help develop clinically useful biomarkers of treatment response to targeted therapies such as TNFi. Mechanism-based patient stratification tools to identify patients likely to respond to treatment may avoid delay in achieving optimal treatment response, prevent the progression of disease and reduce adverse events. Ultimately, these findings may contribute to lowering the cost of treatment and improving patient outcomes in this common chronic inflammatory disease.

## Materials and Methods

### Patients, controls, and PBMC samples

Clinical and biological disease characteristics for the all subjects are summarized in Table 1. Cohort 1 included 48 adults with RA, with PBMC samples collected and cryopreserved at North Shore Long Island Jewish Health System. Controls consisted of 20 healthy age and sex-matched adults at Stanford University School of Medicine. A second group of RA patients was from the NIH-funded Treatment Efficacy and Toxicity in Rheumatoid Arthritis Database and Repository (TETRAD) study, which enrolled 200 RA patients from nine academic medical centers in the US. Inclusion criteria were: age > 19 years; fulfillment of the 1987 ACR classification criteria for RA; and initiating one of the following DMARD or biologic therapies: methotrexate; TNF inhibitors adalimumab, etanercept, golimumab, or infliximab; abatacept; rituximab; or tocilizumab. TETRAD was an observational study, so all medications were started for clinically indicated reasons, and at the direction of the rheumatology provider. High quality, prospectively collected clinical data on disease activity and treatment response were collected from each patient, as were blood samples. Of the 200 TETRAD patients, 181 (90.5%) had high quality cryopreserved PBMCs before and after initiation of the index drug (*13, 20*). Of the 181 RA patients who provided baseline PBMC samples, 146 patients’ samples were included in the baseline pre-treatment analyses (TT0 samples) and 98 patients’ samples were included in the follow up analyses 6 months after initiating index treatment (T6M samples). Patient samples were collected under study protocols approved by the local IRBs at each institution. In accordance with the Declaration of Helsinki, all patients provided written informed consent for the collection and use of their blood samples for research purposes. PMBCs were isolated and cryopreserved as described previously (*37*). A subset (n=11) of healthy controls provided samples at 2 time points one month apart; the induced signaling was compared between the longitudinal samples.

### Single cell network profiling (SCNP) of immune cells from RA and healthy controls

All studies followed the general experimental methods described previously (*37-39*). We designated a stimulus and readout as a node. For example, studying IFNα stimulation within a cell subset using readout of phosphorylation of STAT1 is designated as “IFNα→p-STAT1”. SCNP of 15 nodes (modulator→readout) within 6 immune cell subsets were assayed in all three cohorts (dark blue, Figure 1); additional nodes were measured in TT0 and T6M for a total of 42 and 32 signaling nodes, respectively, in 21 cell subsets (Figure 1). Cryopreserved PBMC samples were thawed, washed, resuspended in RPMI 1640 (10% FBS), aliquoted to 96-well plates at 1 million cells (Cohort 1) and 100,000 cells (TT0, T6M) per well and rested for 2 h at 37°C prior to stimulation with cytokines, TLR agonists, and B cell receptor and T cell receptor crosslinking (Supplemental Table 2). Sample viability was confirmed for all samples by propidium iodide (PI) staining (Cohort 1) or Amine Aqua viability dye (TT0, T6M). Not all TETRAD samples had enough cells for all the conditions to be tested; as a result the conditions were ranked so that biology of most interest was interrogated with more RA samples. Following stimulation for either 2 to 30 minutes depending upon the biology assayed, the cells were fixed with PFA, permeabilized with ice-cold methanol, and stored at −80°C as previously reported (*40*). Cohort 1 modulated conditions were molecularly barcoded with amine-reactive Pacific Orange succinimidyl ester and Alexa 750 carboxylic acid, succinimidyl ester (Invitrogen) and pooled prior to antibody staining, as previously described (*41, 42*). Cells from all cohorts were washed in FACS Buffer (PBS, 0.5% BSA, 0.05% NaN3), stained with cocktails of fluorochrome–conjugated antibodies (Supplemental Table 3), and acquired on LSR II (Cohort 1) or CANTO II (TT0, T6M) flow cytometers using FACS DIVA software (BD Biosciences). All flow cytometry data were analyzed with FlowJo (TreeStar Software, Ashland, OR) or WinList (Verity House Software, Topsham, ME). Cell populations were delineated as shown in Supplemental Table 4.

### Quality Control

The SCNP assay incorporates a number of standardization procedures and process controls that include instrument standardization and calibration, reagent qualification and quality control testing, consistent sample processing via standard operating procedures, and assay performance monitoring (*37*). For TT0 and T6M sample processing, a cell line control row (consisting of the GDM-1, Ramos, or Jurkat cell lines arrayed according to the biology tested, e.g. Jurkat cells as a control for TCR stimulation) were included on each of the 96-well plates that were processed in this study along with a control PBMC sample run with each batch of samples. The controls were used to monitor the reproducibility of the assay performance. Dead cells and debris were excluded by forward scatter (FSC), side scatter (SSC) for all samples and for TT0 and T6M, the additional staining of cleaved PARP, a marker of early apoptosis, was used to gate out dying cells. Propidium iodide staining of Cohort 1 samples confirmed the health of the samples with less than 5% of cells staining positive. We included only node:cell subset that changed in response to stimulation meeting a minimum threshold of log2Fold ≥ 0.3 in the statistical analyses.

### SCNP assay metrics

The “metric” is the quantitative evaluation of signaling protein response, as described previously (*14, 15*). In this study, we used the metric log2Fold to assess the magnitude of response of stimulated cells relative to reference cells. MFIs were obtained for all samples; and for TT0 and T6M samples, the raw data were converted to plate-calibrated ERFs using rainbow calibration particles (Spherotech). For a given cell population and readout, the log2Fold metric measures the magnitude of the responsiveness compared to unstimulated cells using the formula: log2 (ERF_stimulated_/ERF_unstimulated_) where a zero indicates no induced signaling and positive values indicate increases in signaling.

### Data analysis

The two-sample Wilcoxon’s rank sum test was used to compare the phosphorylation of signaling proteins between healthy controls and RA patients for all cohorts and for T6M samples compared to TT0 samples, and the P values <0.05 was considered as statistically significant. Longitudinal samples from healthy controls were analyzed by linear regression and by a variability ratio that assesses assay and biological variability. The variability ratio is determined by: 1) subtracting the signaling of time point 2 from time point 1 to calculate differences and then calculating the median absolute deviation (MAD) of the differences (robust calculation of the standard deviation); 2) calculating the mean signaling for both time points; and then 3) calculating the variability ratio by dividing step 1 by step 2. This statistic is similar to a coefficient of variation (CV). For example, a ratio of 0.1 represents average difference between time points is 10% of mean signal. Signaling associations with RA disease activity (DAS28) and treatment response (EULAR) were identified using linear regression (DAS28) or logistic regression (EULAR). For disease activity, the form of the regression equation is: DAS28 ∼ SCNP+Age. For treatment response, responders (good or moderate) were coded as ‘1’ and non-responders (no response) were coded as ‘0’ for the logistic regression analysis. The form of this regression equation is: Response vs. no response ∼ SCNP. Nodes were considered to be significant if the p-value for the one-sided t-test of the regression coefficient was < 0.05, as calculated using the linear model function “lm” in R. Multivariate models of TNFi response were generated by unsupervised clustering, decision tree analysis, and the Least Absolute Shrinkage and Selection Operator (LASSO) regression method. Performance of models were evaluated through sampling the dataset with replacement 500 times (bootstrap) with ∼2/3 of bootstrap sample used for training and ∼1/3 for testing. Models were evaluated with median model AUROC across bootstraps to determine how well the two groups are discriminated by the models.

### Outcome Measures

We applied SCNP to samples from Cohort 1, TT0 and T6M to dissect the association of signaling pathways with the Disease Activity Score on 28 joints (DAS28) as a measure of disease activity. We used TT0 and T6M samples to explore longitudinal changes and response to TNF inhibitors using the DAS-based European League of Associations against Rheumatism (EULAR) response criteria (*43*). In TETRAD, DAS28 was measured at baseline (TT0) and follow up (6 months after initiation of treatment). Swollen joints and tender joints were scored using 28 version of simplification of original 44 joints score. Blood inflammation marker, erythrocyte sedimentation rate (ESR), was measured by a blood test. We used the following equation (DAS28 with 3 variables) to calculate the Modified Disease Activity Score (DAS): DAS28= (0.56*sqrt (TENDER) + 0.28*sqrt (SWELL) + 0.70*ln (ESR))*1.08 + 0.16. A DAS28 of greater than 5.1 indicates high, 3.2 < DAS ≤ 5.1 moderate, and DAS ≤ 3.2 low disease activity. Changes in DAS28 from before (TT0) to after (T6M) initiation of TNFi treatment (ΔDAS28) were used to classify patients’ responses as good (ΔDAS28 >1.2 and post-treatment DAS28 ≤ 3.2), moderate (0.6<ΔDAS28 ≤1.2 and post-treatment DAS28 < 5.1) or none (ΔDAS28 <0.6).

## Supplementary Materials

**Supplemental Figure 1.** Immune signaling is a stable phenotype in healthy controls.

**Supplemental Figure 2**. Box and whisker plots of modulated signaling (log2Fold) in 6 immune subsets from healthy controls and RA patients from Cohort 1 and T6M.

**Supplemental Table 1.** Baseline Characteristics of 46 TETRAD RA Patients without previous biologic therapy, starting a TNF inhibitor, with 3 month EULAR Response Data and sufficient SCNP data for univariate analysis.

**Supplemental Table 2.** Modulation conditions tested across samples/cohorts.

**Supplemental Table 3.** Antibodies used in these studies.

**Supplemental Table 4.** Phenotypic definitions of cell populations analyzed in these studies.

## Acknowledgments

The authors thank all of the patients with RA and healthy controls who provided samples for this study. This manuscript is dedicated to the memory of Betty Hawtin.

## Data and materials availability

All data associated with this study are present in the paper and/or the Supplementary Materials.

