## Supplemental for "A Systems Immunology Approach Identifies Cytokine-Induced STAT Signaling Pathways Critical to Rheumatoid Arthritis Disease Activity and Treatment Response"

**Supplementary Materials:**

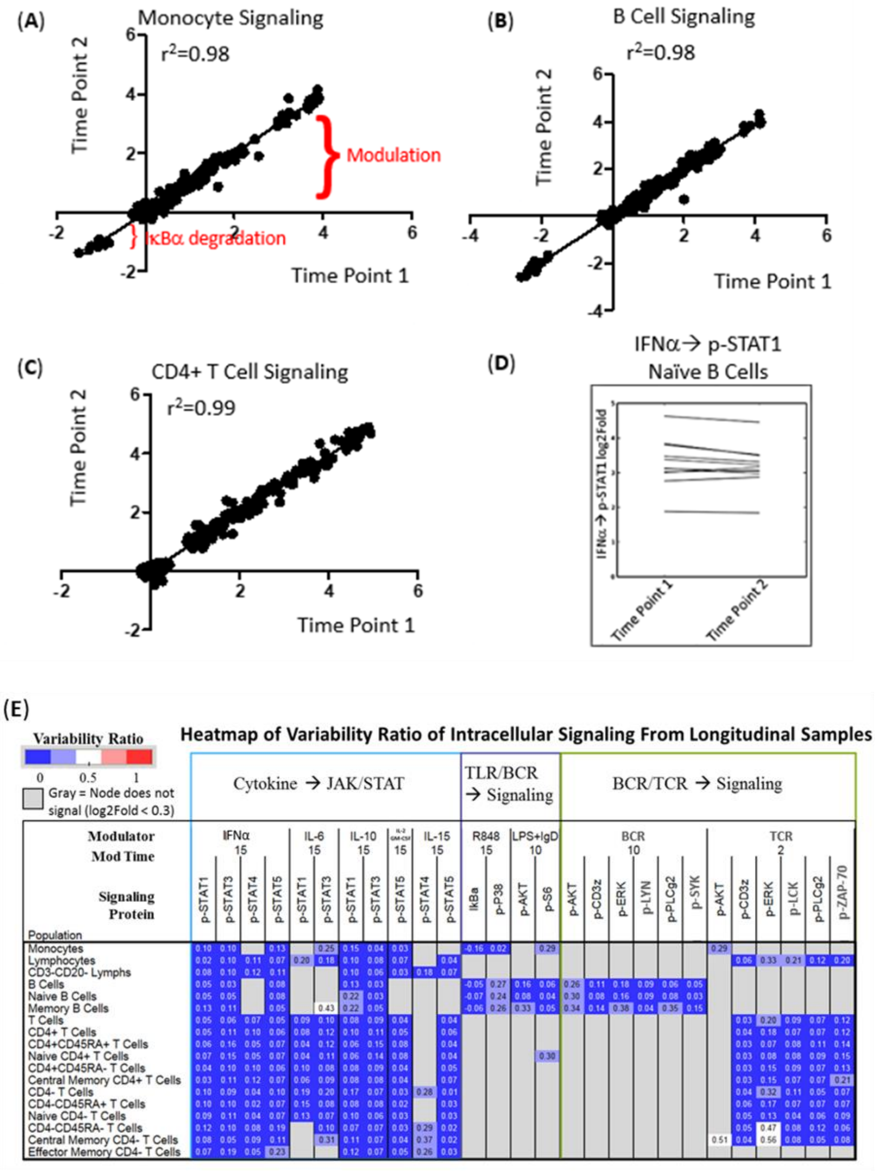


**Supplemental Figure 1.** Immune signaling is a stable phenotype in healthy controls. Monocytes (A), B cells (B), and CD4+ T cells (C) collected from 11 healthy controls at two time points, one month apart, show similar signaling magnitude and patterns. The scale of the log2Fold data ranges from a negative value for IκBα degradation following stimulation that activates NF-kB, to positive values indicating increases in signaling. (D) Line plot shows stable IFNα→p-STAT1 signaling in naive B cells at two time points in 11 healthy controls. (E) Analysis of signaling stability by the variability ratio (see Methods for details); numbers in each cell of the heat map represent the variability of signaling within that node: cell population. This metric is similar to a coefficient of variation (SD/mean), and accounts for assay variability, biological variability, and the magnitude of signaling.

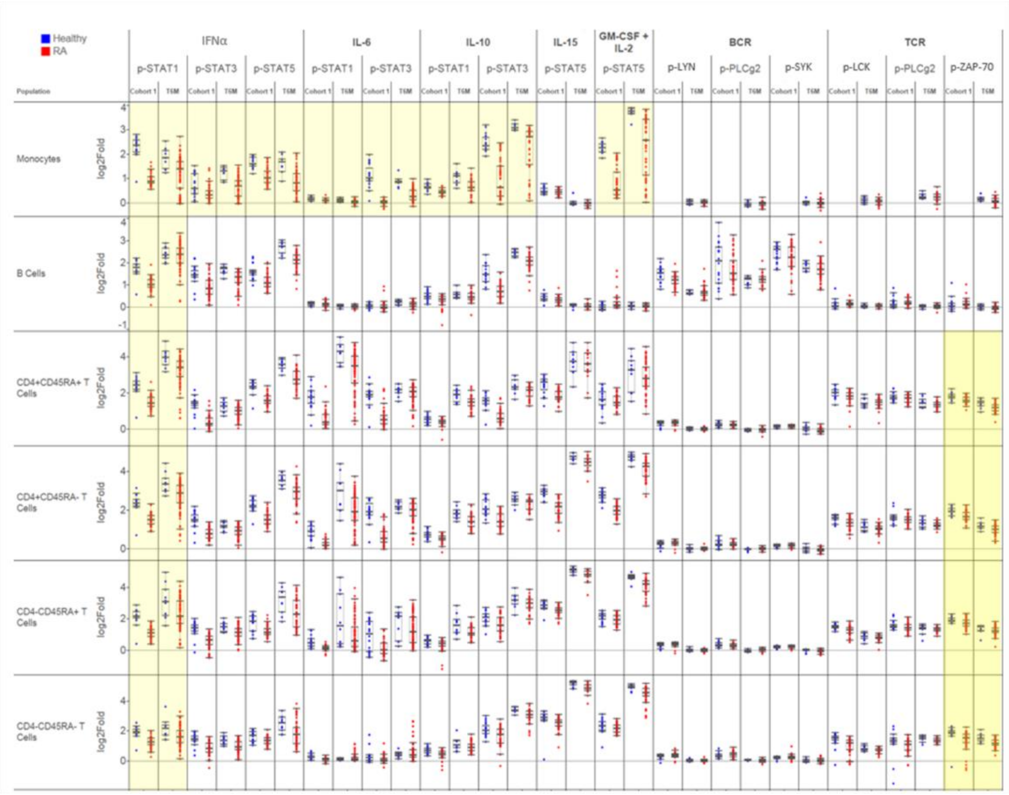


**Supplemental Figure 2.** Box and whisker plots of modulated signaling (log2Fold) in 6 immune subsets from healthy controls and RA patients from Cohort 1 and T6M. Analyses shaded in yellow are shown in detail in Figure 3.

| | | |
|---|---|---|
| Age (median and range) | 55.5 years (25-82) | |
| Sex (no. and % female) | 39 (85%) | |
| Index Drug | Adalimumab | 17 (37%) |
| | Etanercept | 23 (50%) |
| | Golimumab | 2 (4%) |
| | Infliximab | 4 (9%) |
| 3 Month EULAR Response | Good | 9 (20%) |
| | Moderate | 20 (43%) |
| | None | 17 (37%) |
| Autoantibody status (RF or anti-CCP) | Positive | 36 (78%) |
| | Negative | 8 (18%) |
| | Unknown | 2 (4%) |
| Race (self-declared) | African-American | 5 (11%) |
| | Asian | 4 (9%) |
| | Caucasian | 35 (76%) |
| | Other | 2 (4%) |
| Baseline (pretreatment) DAS28 (median and range) | 5.47 (1.97 – 8.22) | |

**Supplemental Table 1.** Baseline Characteristics of 46 TETRAD RA Patients without previous biologic therapy, starting a TNF inhibitor, with 3 month EULAR Response Data and sufficient SCNP data for univariate analysis.

| Modulator | Cohort 1 | TT0 & T6M |
|---|---|---|
| anti-IgD | - | 5 μg/mL (BD) |
| BCR modulation | 10 μg/mL each of anti-IgG (BD), anti-IgM (BD) | 20 μg/mL anti-IgM (Southern Biotech) |
| CD40L | - | 0.5 μg/mL (R&D) |
| Flagellin | - | 10 μg/mL (Invitrogen) |
| GM-CSF | 50 ng/mL (Peprotech) | 10 ng/mL (BD) |
| IFNα | 10,000 IU/mL (Sigma) | 1,000 IU/mL (PBL Assay Science) |
| IL-10 | 50 ng/mL (Peprotech) | 50 ng/mL (BD) |
| IL-15 | 50 ng/mL (Peprotech) | 50 ng/mL (Peprotech) |
| IL-2 | 0.2 ng/mL and 50 ng/mL (BD) | 50 ng/mL (R&D Systems) |
| IL-21 | 50 ng/mL (Peprotech) | 50 ng/mL (Peprotech) |
| IL-6 | 50 ng/mL (BD) | 50 ng/mL (R&D Systems) |
| LPS | - | 1 μg/mL (Sigma) |
| R848 | - | 5 μg/mL (Invivogen) |
| TCR modulation | 10 μg/mL each of anti-CD3ε (BD), anti-mouse (Santa Cruz Biotechnology) | 3 μg/mL anti-CD3ε (Bioscience), 10 μg/mL anti-mouse (Santa Cruz Biotechnology) |
| TNFα | - | 100 ng/mL (BD) |

**Supplemental Table 2.** Modulation conditions tested across samples/cohorts.

| Antibody | Cohort 1 Clone (Vendor) | TT0 and T6M Clone (Vendor) |
| --- | --- | --- |
| CD14 | - | RMO52 (Beckman Coulter) |
| CD19 | HIB19 (eBioscience) | HIB19 (BD) |
| CD20 | - | H1 (BD) |
| CD27 | - | L128 (BD) |
| CD3 | UCHT1 (Invitrogen), UCHT1 (BD) | UCHT1 (BD) |
| CD33 | P67.6 (BD) | - |
| CD4 | RPA-T4 (BD) | RPA-T4 (BD) |
| CD45RA | HI100 (eBioscience) | HI100 (BD) |
| cleaved PARP (D214) | - | F21-852 (BD) |
| IgD | - | IA6-2 (BD) |
| IgM | - | G20-127 (BD) |
| IkB | - | L35A5 (Cell Signaling Technologies) |
| p-AKT(S473) | - | 193H12 (Cell Signaling Technologies) |
| p-CD3ζ(Y142) | - | K25-407.69 (BD) |
| p-ERK(T202/Y204) | - | D13.14.4E (Cell Signaling Technologies) |
| p-LCK(Y505) | 4/LCK-Y505 (BD) | 4/LCK-Y505 (BD) |
| p-p38(T180/Y182) | - | 36/p38 (pT180/pY182) (BD) |
| p-PLCgII (Y759) | K86-689.37 (BD) | K86-689.37 (BD) |
| p-S6 | - | 2F9 (Cell Signaling Technologies) |
| p-STAT1(Y701) | 4a (BD) | 4a (BD) |
| p-STAT3(Y705) | 4/P-STAT3 (BD) | 4/P-STAT3 (BD) |
| p-STAT4(Y693) | - | 38/p-Stat4 (BD) |
| p-STAT5 | 47/Stat5(pY694) (BD) | 47/Stat5(pY694) (BD) |
| p-Zap70(Y319)/p-Syk(Y352) | 17A/P-ZAP70 (BD) | 17A/P-ZAP70 (BD) |

**Supplemental Table 3.** Antibodies used in these studies.

| Population | Definition | Cohort 1 | TT0, T6M |
|---|---|---|---|
| Monocytes | CD14+ or CD33+, high SSC | x | x |
| Lymphocytes | CD14- or CD33-, low SSC | x | x |
| B cells | CD19+ or CD20+ lymphocyte | x | x |
| Naïve B cells | CD20+CD27- lymphocyte | - | x |
| Memory B cells | CD20+CD27+ lymphocyte | - | x |
| T cells | CD3+ lymphocyte | x | x |
| CD4+ T cells | CD3+ CD4+ lymphocyte | x | x |
| CD4+CD45RA+ T cells | CD3+ CD4+ CD45RA+ lymphocyte | x | x |
| Effector CD4+ T cells | CD3+ CD4+ CD45RA+ CD27-lymphocyte | - | x |
| Naive CD4+ T cells | CD3+ CD4+ CD45RA+ CD27+ lymphocyte | - | x |
| CD4+CD45RA- T cells | CD3+ CD4+ CD45RA- lymphocyte | x | x |
| Effector Memory CD4+ T cells | CD3+ CD4+ CD45RA- CD27- lymphocyte | - | x |
| Central Memory CD4+ T cells | CD3+ CD4+ CD45RA- CD27+ lymphocyte | - | x |
| CD4- T cells | CD3+CD4- lymphocyte | x | x |
| CD4-CD45RA+ T cells | CD3+ CD4-CD45RA+ lymphocyte | x | x |
| Effector CD4- T cells | CD3+ CD4- CD45RA+ CD27-lymphocyte | - | x |
| Naive CD4- T cells | CD3+ CD4- CD45RA+ CD27+ lymphocyte | - | x |
| CD4-CD45RA- T cells | CD3+ CD4- CD45RA- lymphocyte | x | x |
| Effector Memory CD4- T cells | CD3+ CD4- CD45RA- CD27- lymphocyte | - | x |
| Central Memory CD4- T cells | CD3+ CD4- CD45RA- CD27+ lymphocyte | - | x |
| CD3-CD20- Lymphs (NK enriched) | CD3-CD20- lymphocyte | x | x |

**Supplemental Table 4.** Phenotypic definitions of cell populations analyzed in these studies.
